# BRAF inhibitor resistance confers increased sensitivity to mitotic inhibitors

**DOI:** 10.1101/2021.04.15.439990

**Authors:** SA Misek, TS Dexheimer, M Akram, SE Conrad, JC Schmidt, RR Neubig, KA Gallo

## Abstract

Single agent and combination therapy with BRAF^V600E/K^ and MEK inhibitors have remarkable efficacy against melanoma tumors with activating BRAF mutations, but in most cases resistance eventually develops. The purpose of this study is to uncover pharmacological vulnerabilities of BRAFi-resistant melanoma cells, with the goal of identifying new therapeutic options for patients whose tumors have developed resistance to BRAFi/MEKi therapy. We screened a well-annotated compound library against a panel of isogenic pairs of parental and BRAFi-resistant melanoma cell lines to identify classes of compounds that selectively target BRAFi-resistant cells over their BRAFi-sensitive counterparts. Two distinct patterns of increased sensitivity to classes of pharmacological inhibitors emerged. In two cell line pairs, BRAFi resistance conferred increased sensitivity to compounds that share the property of cell cycle arrest at M-phase, including inhibitors of aurora kinase (AURK), polo-like kinase (PLK), tubulin, and kinesin. Live cell microscopy used to track mitosis in real time revealed that parental, but not BRAFi-resistant, melanoma cells were able to exit from compound-induced mitotic arrest through mitotic slippage, thus escaping death. Consistent with the key role of Cyclin B1 levels in regulating mitosis at the spindle checkpoint, in arrested cells we found higher Cyclin B1 levels in parental over BRAFi-resistant melanoma cells, suggesting that altered Cyclin B1 expression levels may explain why these BRAFi resistant cells have gained increased vulnerability to mitotic inhibitors. Another BRAFi-resistant cell line showed increased sensitivity to Chk1/2 inhibitors, possibly due to an accumulation of DNA damage, resulting in mitotic failure. This study shows that BRAFi-resistance in melanoma cells confers vulnerability to pharmacological disruption of mitosis and suggests a targeted synthetic lethal approach to treat BRAF-mutant melanomas that have become resistant to BRAF/MEK-directed therapies.

## Introduction

Many mechanisms of BRAFi/MEKi resistance in BRAF-mutant melanoma are well understood (*1-12*), yet systematic approaches to identifying effective second-line therapies are still largely lacking. One appealing strategy to treat drug-resistant melanoma is to re-purpose drugs that have been FDA-approved for other indications since they can be quickly translated to the clinic. Large-scale efforts have sought to systematically profile compounds against annotated panels of cancer cell lines, initially with datasets like Genomics of Drug Sensitivity in Cancer (GDSC) (*13*) or Cancer Target Discovery and Development (CTD^2^) (*14*), and more recently with Profiling Relative Inhibition Simultaneously in Mixtures (PRISM) (*15, 16*). The ultimate goal of each of these initiatives is to correlate genomic features with drug responses and map those associations back to patient tumors. Additional targeted screens have also been used to identify compounds with activity against drug-resistant cancer models (*17-20*).

The strategy we implemented was to screen a library of chemical compounds against pairs of isogenic parental and BRAFi-resistant melanoma cell lines. Chemical compound screens compare well with functional genomics-based CRISPR screens, but also present several distinct advantages. Most standard CRISPR screens are based upon perturbation of individual genes often leading to compensation by redundant isoforms, whereas compound screens typically contain inhibitors that target multiple members of the same protein family. Furthermore, CRISPR screens typically rely on measurement of responses that require long-term deletion of a target gene. Thus, if a gene is essential for survival of all cells, it is impossible to assess the differential dependence of various cell populations on that gene. Finally, a drug repurposing approach immediately highlights promising drug candidates with activity against the target cells that could be translated to the clinic.

Mitotic inhibitors are often effective therapies for treating cancer as they induce mitotic arrest, followed by cell death. However, resistance to antimitotic therapies can occur when cancer cells undergo mitotic slippage, allowing them to them to survive in a polyploid state. Cyclin B1 levels are critical in regulating cell cycle exit from mitotic arrest. Gradual degradation of Cyclin B1 during prolonged cell cycle arrest results in premature chromosome decondensation (*21*) and cells subsequently exit from the cell cycle into a 4n state. These cells are senescent, but under certain conditions such as loss of p53 they can re-enter into the cell cycle (*22*). Since mitotic slippage initially gives rise to tetraploid cells, subsequent rounds of mitosis in cells which have undergone mitotic slippage will give rise to polyploid cells.

In this study, our screen which was designed to reveal pharmacological vulnerabilities of BRAFi-resistant melanoma cells, identified compounds that disrupt mitosis through multiple, distinct mechanisms. For instance, Aurora kinase (AURK) and Polo-like kinase (PLK) inhibitors as well as inhibitors of tubulin polymerization arrest cells in mitosis and prevent chromosome alignment during metaphase. These classes of compounds selectively induce prolonged cell cycle arrest and apoptosis in BRAFi-resistant cells. In contrast we found that the parental melanoma cells were markedly less sensitive to mitotic inhibitors.

We further elucidated the mechanistic basis for this selectivity by demonstrating that after treatment with an AURK inhibitor parental melanoma cells have a greater propensity to undergo mitotic slippage than their BRAFi resistant counterparts. We observed lower levels of Cyclin B1 in parental compared with BRAFi resistant melanoma cells after treatment with AURK inhibitor. These findings are consistent with the model in which parental BRAF mutant melanoma cell lines retain the ability to degrade Cyclin B1 and thus evade mitotic inhibitor-induced death by undergoing mitotic slippage, whereas their BRAFi resistant counterparts are unable to downregulate Cyclin B1 and thus undergo mitotic inhibitor induced death. Not all BRAFi resistant melanoma lines share this enhanced selectivity for mitotic inhibitor. Our screen revealed that one BRAFi-resistant melanoma cell line is more sensitive to pharmacological inhibition of Chk1/2 than its isogenic parental cell line. We hypothesize this is due to accumulation of DNA damage which results in mitotic failure, and ultimately cell death. In summary our work has identified new potential approaches to treating BRAFi resistant melanomas. Furthermore, we have probed two distinct mechanisms through which BRAFi-resistance in melanoma cells leads confers new vulnerability to pharmacological disruption of mitosis. These studies open up the exciting possibility that mitotic inhibitors may serve as potential new treatment strategies for BRAFi-resistant melanoma tumors. In addition, exploiting these vulnerabilities may be valuable in preventing the development of BRAFi resistance outright.

## Materials and Methods

### Cell lines, reagents, and antibodies

Parental (denoted by a P suffix in the cell line name) and matched isogenic BRAFi-resistant cells (denoted by an R suffix in the cell line name) were either a gift from Dr. Roger Lo (UCLA) (M229P/R, M238P/R, or M249P/R)(*7*) or generated in our laboratory (UACC62P/R), as previously described (*23*).

BI-2536 (#17385), Volasertib (#18193), GSK461364 (#18099), Danusertib (#18387), AMG900 (#19176), MLN8237 (#13602), Docetaxel (#11637), Ispinesib (#18014), Mebendazole (#18872), AZD7762 (#11491), LY2603618 (#20351), SCH900776 (#18131), and Vemurafenib (#10618) were purchased from Cayman Chemical (Ann Arbor, USA). All compounds were diluted in DMSO to a stock concentration of 10 mM and aliquots were stored at -20°C. An antibody against γH2AX (#9718) was purchased from Cell Signaling Technology (Danvers, USA). Alexa Fluor goat anti-rabbit488 (#A11034) was purchased from Invitrogen (Carlsbad, USA). Recombinant human TNFα protein (#210-TA-005) was purchased from R&D Systems (Minneapolis, USA).

### Cell culture

Cells were cultured in DMEM (ThermoFisher, Waltham, USA #11995-065) supplemented with 10% fetal bovine serum (FBS) (ThermoFisher, #10437-028) and 1% Antibiotic-Antimycotic (ThermoFisher, #15240062) and were passaged at approximately 75% confluence. The BRAFi-resistant cell line variants were maintained in culture medium supplemented with 2 µM vemurafenib. Vemurafenib was removed from the culture medium when cells were seeded for experiments, except where otherwise indicated. Cells were routinely tested for mycoplasma contamination by DAPI staining. Short Tandem Repeat profiling of all cell lines was performed at the MSU genomics core. In all cases, isogenic pairs of cell lines had identical STR profiles.

### Generation of recombinant constructs

Scarlet-H2A was amplified using PCR (donor plasmid: Addgene #85051, from Dorus Gadella) and subcloned into pDONR221 using the Gateway BP Clonase II enzyme mix (#11789020) from ThermoFisher. It was subsequently subcloned into the pLX301 lentiviral expression vector (from David Root, Addgene plasmid #25895) using the Gateway LR Clonase II enzyme mix (#11791020) from ThermoFisher. TUBA1B was amplified using PCR (donor plasmid: Addgene #57159, from Michael Davidson) and an EGFP-TUBA1B fusion protein was generated with two-stage overhang extension PCR using the TUBA1B and EGFP cDNA fragments. The EGFP-TUBA1B fusion protein was subcloned into pDONR221 and was subsequently cloned into pLX303 (from David Root, Addgene #25897). CyclinB1-GFP was amplified using PCR (donor plasmid: Addgene #26061, from Jonathon Pines) and was subcloned into pDONR221 and subsequently subcloned into pLX303. All PCR primers are listed in Table S1. Successful cloning was confirmed by Sanger sequencing.

### Virus generation and infection

HEK-293T cells were seeded onto 10-cm plates at a density of 4×10^6^ cells/plate and the cells were allowed to attach overnight. The next day the cells were transfected with a plasmid cocktail containing 5000 ng of the pLentiCRISPRv2 plasmid, 5000 ng of psPAX2 (Addgene plasmid #12260), 500 ng of pMD2.G (Addgene plasmid #12259), and 20 µL of Lipofectamine 2000 (ThermoFisher, #11668019) in 400 µL of OptiMEM (ThermoFisher, #31985070). The next morning the medium was changed to 10 mL of fresh complete culture medium, and the following day each plate was supplemented with an additional 5 mL of culture medium. After 24 h, the culture medium was harvested and filtered through a 0.45-µm syringe filter. Virus was stored at 4°C and used within 2 weeks.

Melanoma cells were seeded onto 10-cm plates at a density of 5×10^5^ cells/plate in 10 mL of complete culture medium. Prior to adherence of cells, 3 mL of viral supernatant was added to each plate. The cells were incubated with virus for 24 h, then the medium was changed to 10 mL of fresh medium. After at least 7 days the cells were used in live cell imaging experiments.

### Viability experiments

Cells were seeded into white 384-well tissue culture plates (PerkinElmer, Waltham, USA, #6007689) at a density of 1000 cells/well in 20 µL of growth medium. The next day, compounds were pre-diluted in growth medium and then added to the 384-well plates so that the final volume of each well was 40 µL. A PBS or growth medium barrier was added to the outer wells of the plate to limit evaporation. Cells were cultured under these conditions for 72 h. To assess viability, 8 µL of CellTiter-Glo (Promega, Madison, USA, #G7573) was added to each well. Plates were incubated on an orbital shaker for 5 min at room temperature, then briefly centrifuged (4000 rpm, 60 s) before being read on a Bio-Tek Synergy Neo plate reader with the #11 and #41 Ex/Em filter cubes. Viability signal was plotted versus log (vemurafenib concentration) for each treatment condition.

### Compound library screen

Cells were seeded into white 384-well plates at a density of 1,000 cells/well. The next day the NCATS MIPE chemical library (*24*) was pinned into the plates at a final concentration of 200 nM. After 72 h, 8 µL of CellTiter-Glo was added to each well. The plates were incubated on an orbital shaker for 5 min, briefly spun down, and cell viability was measured as described above. In some cases, noise in the assay produced viability measurements that were greater than 100%. In these situations, the viability measurement was set to 100%. Viability data from the compound screen is listed in Table S2.

### Cell cycle analysis

Cells were rinsed once in PBS, incubated with trypsin, washed once in PBS and immediately fixed in 70% ethanol for 20 min at room temperature. The cells were washed once and were re-suspended in PBS supplemented with 20 µg/mL propidium iodide (#P1304MP, ThermoFisher) and 200 µg/mL RNaseA. The cells were briefly mixed and were incubated on ice for 20 min. Following incubation, the cells were filtered through a 70 µM filter and were run on an Accuri C6 flow cytometer (BD Biosciences, Franklin Lakes, USA). Data were analyzed with the FCS Express flow cytometry analysis software package.

### Assay for reactive oxygen species

Cells were seeded at a density of 10,000 cells/well in a 96-well plate and allowed to attach overnight. The next day reactive oxygen species (ROS) levels were measured. Cells were also treated with 1 mM H_2_O_2_ for 15 min as a positive control. The ROS assay (#MAK145, Sigma-Aldrich, St. Louis, USA) was performed as described in the manufacturer’s protocol for adherent cells.

### Immunofluorescence staining

Cells were seeded into 8-well chamber slides and were treated as indicated in the figure legends. Cells were fixed with 3.7% formaldehyde for 15 min then blocked in 2% BSA PBS-Triton X-100 (0.1%) for 1 h at room temperature. Cells were incubated overnight at 4°C in phospho-γH2AX antibody at a dilution of 1:1,000 in blocking buffer. Cells were washed thrice in PBS and then were incubated in the appropriate secondary antibody at a 1:1,000 dilution for 1 h at room temperature. Cells were washed 3 times in PBS and slides were then mounted in ProLong Gold Antifade + DAPI (ThermoFisher, #P36935). Slides were cured overnight at room temperature, and then transferred to 4°C. Slides were imaged on a Nikon TE2000-U fluorescence microscope at 20x magnification. All images were automatically quantified using an ImageJ pipeline. Briefly, nuclear masks were created from the DAPI channel and the phospho-γH2AX staining intensity was measured within each mask. Data is reported as relative phospho-γH2AX fluorescence intensity. At least 500 cells were quantified per treatment condition.

### Live cell imaging

To quantify the rate and outcome of mitosis in melanoma cells, UACC62P/R and M229P/R cells were engineered to express Scarlet-H2A and EGFP-TUBA1B. Cells were seeded at a density of 5,000 per well in a glass-bottom 96-well plate. The next day the cells were treated as described in the figure legends and were imaged at 3-min intervals on a BioTek Cytation 3. Over 40 cells per treatment condition were analyzed for mitotic rate and outcome. The T_0_ for mitotic entry was defined as nuclear envelope breakdown and the final time was defined as either completion of mitosis (chromosome segregation and complete de-condensation), mitotic slippage (complete de-condensation of chromosomes), or prolonged arrest at the end of imaging.

To generate high resolution images, cells were seeded at a density of 10,000 per well in 8-well glass-bottom chamber slides. The next day the growth medium was changed to CO_2_-independent growth medium (Gibco, #18045088) and the cells were treated as described in the figure legends. Cells were imaged with a 20x air objective on a DeltaVision microscope equipped with an sCMOS camera, environmental chamber, and ultimate focus drift correction system. Five z-sections were imaged in 2 µm steps at 3-min time intervals. Equivalent exposure conditions were used for all images.

The described DeltaVision setup and imaging parameters were used to generate quantitative Cyclin B1 protein expression data. At least 10 cells were analyzed per treatment condition. Cyclin B1 expression was quantified at each time interval in with FIJI v1.52p. Cyclin B1 expression was normalized to the expression value at the first analyzed timepoint.

## Results

### BRAFi-resistant melanoma cells are sensitive to inhibitors that disrupt mitosis

In this study, we sought to identify compounds that selectively target BRAFi-resistant melanoma cells as potential therapeutic strategies and as a window to understanding mechanisms through which resistance arises. In our initial screen we profiled the NCATS Mechanism Interrogation PlateE (MIPE) library of 1910 compounds (*24*) against a pair of matched isogenic parental and BRAFi-resistant melanoma cells, UACC62P which harbors the BRAF^V600E^ oncogene and its resistant counterpart UACC62R which was developed by *in vitro* selection with vemurafenib (*23*). The NCATS MIPE library contains a mechanistically and structurally diverse set of compounds, the majority of which are either FDA-approved or investigational new drugs and are directed at over 900 unique protein targets. The library is also redundant, containing multiple inhibitors against any one individual protein. This approach allows us to not only identify efficacious compounds, but also to gain new mechanistic insights into the molecular mechanisms of BRAFi resistance. Fig. 1A shows a graphical representation of sensitivity of each compound against the UACC62P (x-axis) and UACC62R (y-axis) cell lines. As expected, unlike the parental counterpart, the UACC62R cells were insensitive to RAF and MEK inhibitors in this library, demonstrating that our screen can identify compounds that differ in their selectivity towards the parental and BRAFi-resistant melanoma cells. As shown in Fig. 1A, compounds that target PLK, AURK, tubulin, and kinesin selectively reduced viability of the UACC62R cells. Since the screen was performed at a single concentration of each compound, fresh powder was used to validate in concentration response studies 9 of the screen hits, which included 3 PLK inhibitors (BI2536, Volasertib, and GSK461364), 3 AURK inhibitors (Danusertib, AMG900, and MLN8237), 2 tubulin inhibitors (Docetaxel and Mebendazole), and the kinesin inhibitor Ispinesib. All of these top hits were validated (Fig. 1B). Interestingly, the differential compound sensitivity was found to be due to a change in the maximum percent inhibition (E_max)_, rather than due to a difference in the IC_50_ (Fig. 1B). Our results indicate that mitotic blockade selectively reduces viability of BRAFi resistant melanoma cells. No obvious synergy was observed between vemurafenib and any of the identified compounds (Fig. S1). This suggests that alterations unique to the UACC62R cells render them more vulnerable to disruption of mitosis.

**Figure 1.**
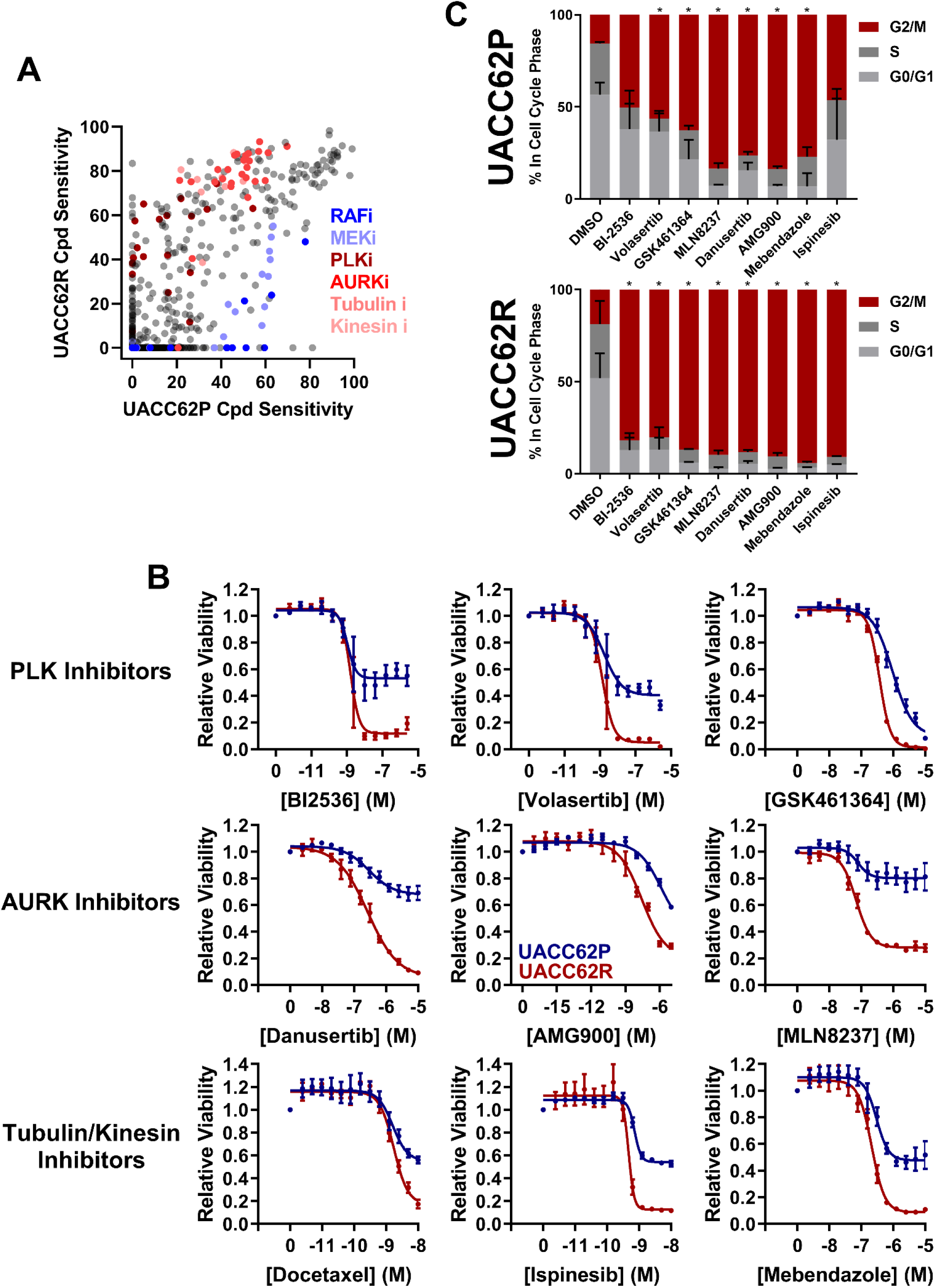
Vemurafenib-resistant UACC62R cells are selectively vulnerable to pharmacological disruption of mitosis. **A**. The NCATS MIPE chemical library was screened against parental and resistant UACC62P/R cells at 200 nM as described in the Materials and Methods section. Compound sensitivity data are plotted as % reduction in viability of UACC62P cells vs UACC62R cells for each compound in the screen. The larger the sensitivity value, the greater was the reduction in cell viability. The screen was performed with n = 1 replicates for each cell line. Inhibitors with selected targets are indicated in shades of *blue* for those that showed greater efficacy in UACC62P cells, and in shades of red for those that showed greater efficacy in UACC62R cells. **B**. Fresh powder for 9 of the compounds identified in the initial screen was obtained and the effect of these compounds on cell viability was analyzed at the indicated concentrations. *Blue* lines represent data for the UACC62P cells, and *red* lines indicate data for UACC62R cells. Data are represented as mean ± SE of the technical replicate averages (n=3) for each of the biological replicates (n = 3). IC_50_ and E_max_ values are listed in Table S3. **C**. Cell cycle analyses of vehicle and drug-treated UACC62P/R cells were performed as described in the Materials and Methods section. All compounds were used at concentrations of 1 µM except for Ispinesib which was analyzed at 1 nM. Statistical analyses were performed on the proportion of cells in G2/M for the drug-treated samples vs the DMSO control using One-way ANOVA analysis, * indicates p < 0.01. Data are represented as mean ± SE for n = 3 biological replicates.

We next expanded the screen to include two additional BRAF^V600E^ vemurafenib-sensitive/vemurafenib-resistant melanoma cell line pairs, M238P/R and M229P/R (*7*) which share similar transcriptional profiles with the UACC62P/R cells. In particular, compared with their vemurafenib-sensitive parental counterparts, the resistant cells lack expression of differentiation-associated melanocyte lineage genes (*23*). M238R cells showed a compound sensitivity pattern similar to UACC62R, with top hits including AURK inhibitors (Fig. S2), whereas M229R cells showed increased sensitivity to Chk1/2 inhibitors over its parental counterpart (Fig. S2). Interestingly, the identified AURK inhibitors that selectively target UACC62R over UACC62P cells were different from those that target M238R over M238P cells (Table. S2). Differences in expression or activities of drug efflux pumps or drug metabolizing enzymes in the various BRAFi resistant melanoma lines could provide an explanation for these findings.

As a control experiment, we also screened the M249P and M249R melanoma pair, which has a transcriptional profile that is distinct from the other vemurafenib-resistant melanoma cells. In M249R cells, vemurafenib resistance has been shown to be due to acquisition of the activating NRAS^Q61^ mutation (*7*) leading to reactivation of the ERK/MAPK pathway. In our screen, there was no enrichment of mitotic inhibitors with selectivity towards M249R cells, consistent with the finding that resistance develops through MAPK reactivation (Fig. S2/Table S2).

PLK, AURK, tubulin, and kinesin are all critical for the execution of mitosis, so we reasoned that altered regulation of mitosis might provide the mechanistic basis for the differences in selectivity between the UACC62P and UACC62R cells. Therefore, we performed cell cycle analysis to determine the impact of the mitotic inhibitors identified in our screen on cell cycle distribution of UACC62P and UACC62R cells. These data show that treatment with mitotic inhibitors results in a greater fraction of UACC62R cells with 4n DNA content (G2/M) compared with UACC62P, indicating that the UACC62R cells undergo more efficient mitotic arrest in response to drug treatment than their parental counterpart (Fig. 1C).

### Parental, but not BRAFi-resistant, melanoma cells undergo mitotic slippage

Our data demonstrate that BRAFi-resistant cells are more sensitive than their parental counterparts to inhibitors which disrupt mitosis. However, the mechanism behind this increased sensitivity was initially unclear. We first hypothesized that there might be increased levels of DNA damage in BRAFi-resistant cells which could enhance their sensitivity to pharmacological disruption of mitosis. However, neither ROS, which could in principle induce DNA damage, nor γH2AX staining, a marker of DNA damage, was elevated in UACC62R cells over levels in UACC62P cells (Fig. S3 and S4). We previously described that, compared with their parental counterparts, UACC62R melanoma cells express genes associated with de-differentiation. To investigate whether the increased sensitivity of UACC62P cells to mitotic inhibitors might be attributed to their more differentiated state compared with UACC62R cells, we treated both UACC62 P and UACC62R cell with tumor necrosis factor α (TNFα) which has been shown to induce de-differentiation of melanoma cells (*25, 26*), and assessed the impact on sensitivity to a panel of mitotic inhibitors (Fig. S5). The lack of effect of TNFα on sensitivity to mitotic inhibitors suggests that the de-differentiation attributes of UACC62R cells do not explain their vulnerability to mitotic inhibitors (Fig. S5).

We then sought to determine how mitosis is affected in BRAFi-resistant and isogenic parental cells treated with or without mitotic inhibitors. Fusion proteins of enhanced green fluorescent protein with the α-tubulin B chain (EGFP-TUBA1B) and of a red fluorescent protein with histone H2A (mScarlet-H2A) were used to label the mitotic spindle and chromosomes, respectively. We hypothesized that mitotic integrity in treatment-naïve UACC62R cells might already be impaired, rendering them more vulnerable to pharmacological disruption of mitosis than the non-resistant parental cells. However, DMSO-treated UACC62P and UACC62R cells had similar mitotic timing duration and success rates (Fig. 2A). In contrast to the differential effects of compound-treatment on cell viability (Fig. 1B), treatment with GSK461364 (PLKi), MLN8237 (AURKi), or Mebendazole (tubulin polymerization inhibitor), almost completely prevented both UACC62P and UACC62R cells from successfully completing mitosis. Interestingly, a significant fraction of the compound-treated UACC62P cells initially arrested in mitosis, but after several hours underwent mitotic slippage (Fig. 2B/C). In contrast very few of the compound-treated UACC62R cells did the same. The proportion of cells that undergo mitotic slippage in response to mitotic inhibitor drug treatment was found to inversely correlate with the drug induced decrease in viability (Fig. 1B) which may explain why mitotic disrupters selectively target UACC62R cells.

**Figure 2.**
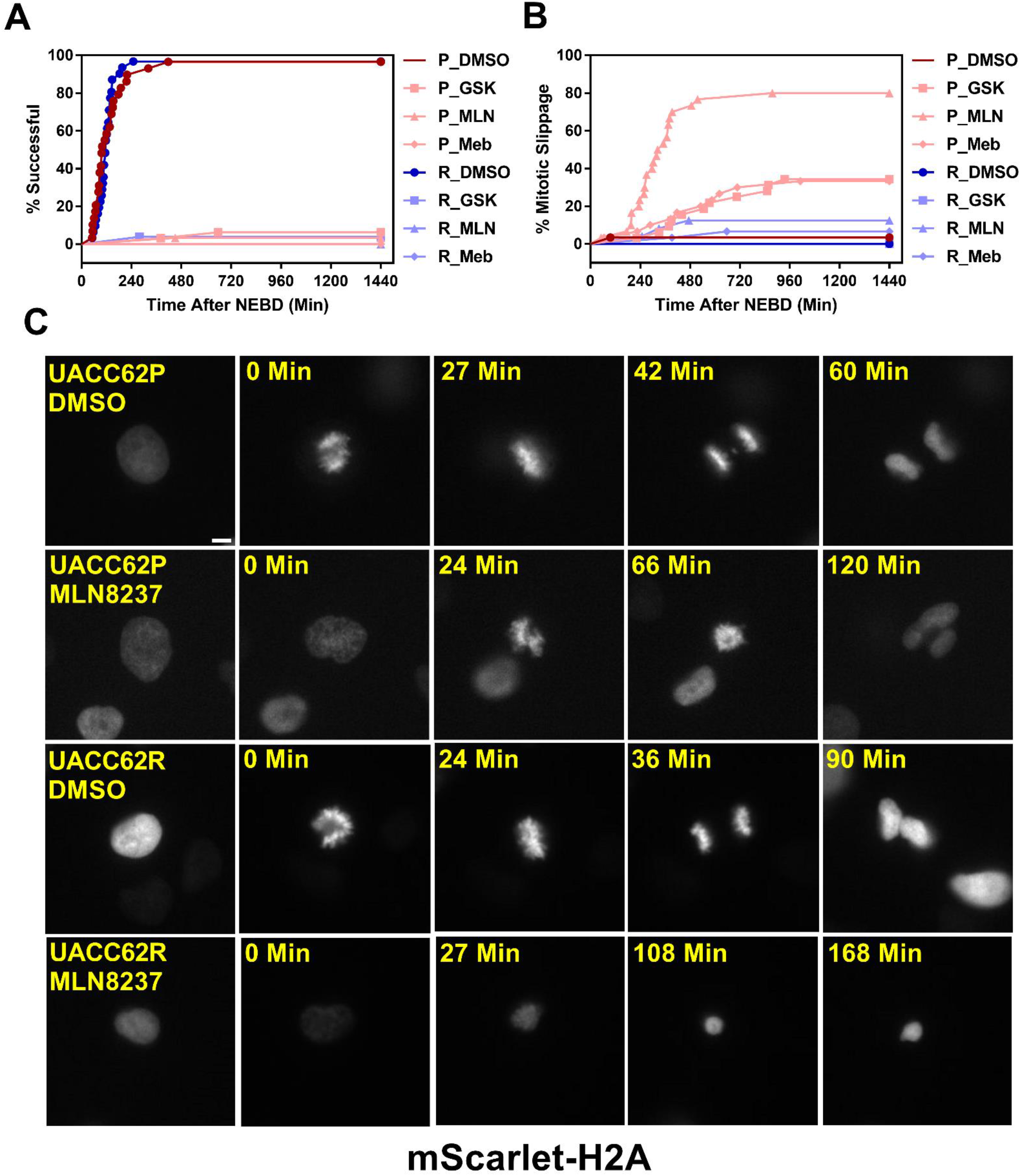
Compound-treated UACC62P, but not UACC62R, cells undergo mitotic slippage. UACC62P/R cells were engineered to stably express GFP-TUBA1B and mScarlet-H2A. The cells were seeded into glass-bottom 96-well plates and the next day the cells were treated with 1 µM GSK461364, MLN8237, or Mebendazole. Mitotic timing and outcomes were analyzed as described in Materials and Methods. The fraction of cells which **A**. successfully completed mitosis or **B**. underwent mitotic slippage are plotted as a function of time. At least 40 cells were analyzed per treatment condition. **C**. Representative images of DMSO or MLN8237-treated UACC62P/R cells. Images were captured using the DeltaVision microscopy setup as described in the Materials and Methods section. Scale bar = 10 µM.

### Differential Cyclin B1 accumulation in UACC62P/R cells

Degradation of Cyclin B1 drives the exit of cells from mitosis. In arrested cells, however, a failure to reduce Cyclin B1 levels below a critical threshold can result in cells undergoing mitotic slippage leading to greater than 2n DNA content and polyploid nuclei (*21*). We therefore hypothesized that our finding that UACC62P cells, but not UACC62R cells, undergo mitotic slippage upon treatment with inhibitors might be due to differences in the levels of Cyclin B1 at the mitotic spindle checkpoint. To explore this idea, we engineered UACC62P/R cells to stably express EGFP-CCNB1 (Cyclin B1) along with mScarlet-H2A to simultaneously monitor in real time both mitotic progression and Cyclin B1 levels by live cell imaging. EGFP-Cyclin B1 expression and localization mirrors that of endogenous Cyclin B1 (*27*) and expression of EGFP-Cyclin B1 does not have a significant effect on cell cycle progression or expression of cell cycle-related genes (*28*). Prior to the initiation of mitosis, EGFP-Cyclin B1 is sequestered in the cytosol in DMSO-treated UACC62P cells and then rapidly co-localizes with mScarlet-H2A upon chromosome condensation and nuclear envelope breakdown (Fig. 3A). The DMSO-treated UACC62R cells displayed kinetics of EGFP-Cyclin B1 expression similar to that of DMSO-treated UACC62P cells (Fig. 3B). In response to treatment with the AURKi, MLN8237, the levels of Cyclin B1 in UACC62R cells gradually reduced to approximately 50% of their original levels. In contrast, in the UACC62P cells treated with MLN8237, there was a greater reduction in the levels of EGFP-Cyclin B1, which could allow these cells to undergo mitotic slippage. Taken together, these data suggest that differential regulation of Cyclin B1 degradation dictate whether MLN8237-treated melanoma cells undergo mitotic slippage or prolonged cell cycle arrest and subsequent loss of viability.

**Figure 3.**
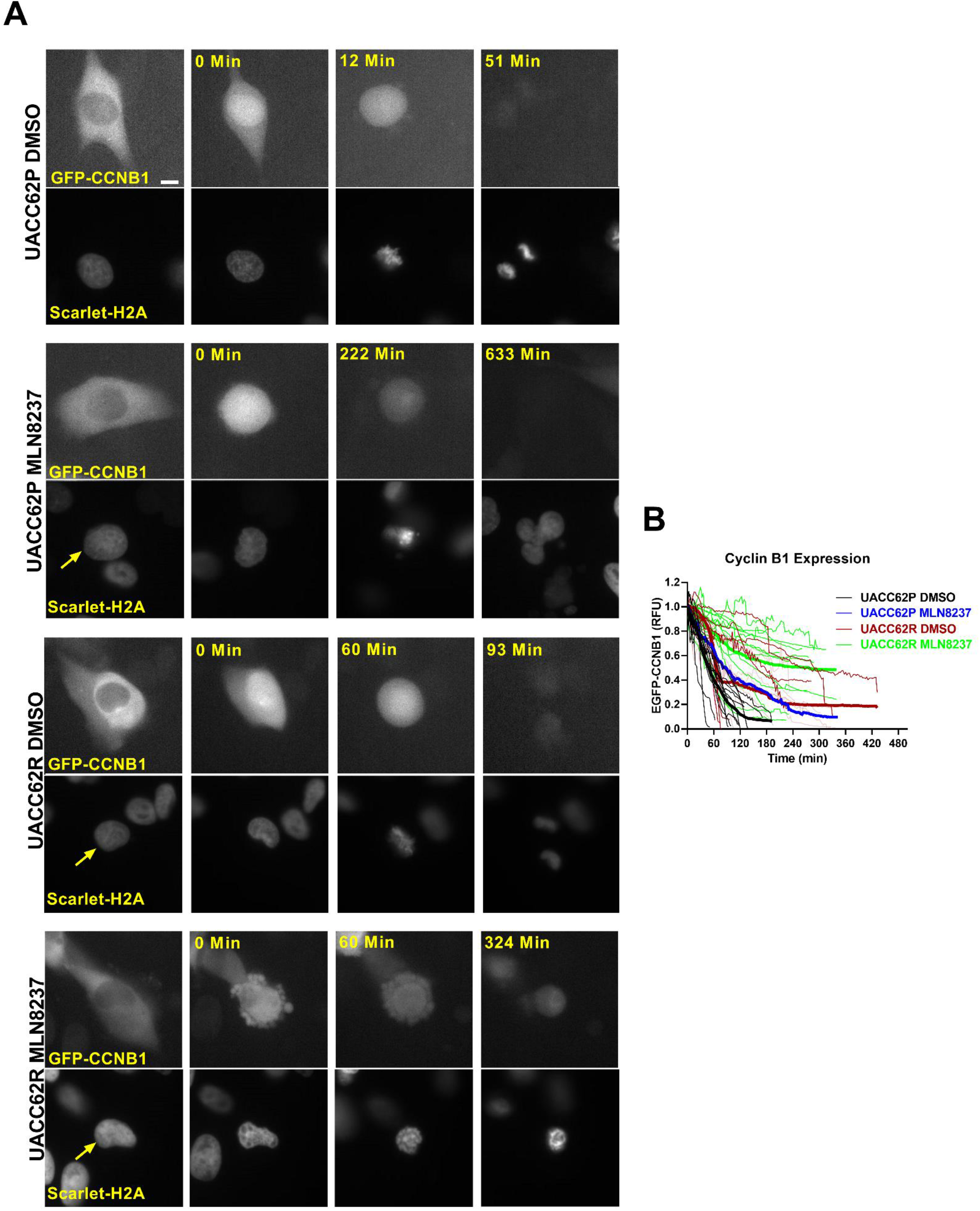
Differential CyclinB1 degradation rates in UACC62P/R cells. **A**. Representative images of EGFP-CyclinB1 and mScarlet-H2A in DMSO or MLN8237-treated UACC62P/R cells. Scale bar = 10 µM. **B**. Quantification of CyclinB1 expression levels in DMSO or MLN8237-treated UACC62P/R cells was performed as described in Materials and Methods. At least 10 cells were analyzed per treatment condition. Thin lines indicate individual cells and bolded lines are the average between all cells in the treatment condition.

### Increased sensitivity of BRAFi-resistant M229R cells to Chk1/2 inhibitors

Our initial compound screen showed that while vemurafenib resistance led to increased sensitivity to inhibitors of AURK, PLK, tubulin, and kinesin in UACC62 and M238 melanoma cells, our screen revealed that M229R cells had increased sensitivity to 3 different Chk1/2 inhibitors compared with M229P cells (Fig. S2). In a follow-up concentration response assay, we confirmed that the three Chk1/2 inhibitors selectively target M229R cells over the vemurafenib sensitive parental cell line (Fig. 4A). Similar to our findings with the AURK/PLK/tubulin/kinesin inhibitors in our previously screened melanoma cells, these inhibitors show no synergy with vemurafenib (Fig. S6). While the mitotic success rate in response to Chk1/2 inhibitor treatment was reduced in M229R cells compared with M229P cells, the fraction of cells undergoing mitotic slippage was identical in M229P and M229R cells (Fig. 4B). After 240 min approximately 70% of compound-treated M229P cells had completed mitosis whereas, depending on the Chk1/2i used, only 30-60% of M229R cells had successfully completed mitosis. These data suggest that while M229R cells are also differentially sensitive to mitotic disrupters, in this case Chk1/2 inhibitors, their increased vulnerability compared to M229P cells appears to be due to a mechanism distinct from mitotic slippage. Under physiological conditions Chk1/2 activation monitors DNA fidelity during replication and in response to DNA damage, ultimately preventing premature entry into mitosis (*29*). Chk1/2 inhibition would be expected to result in the accumulation of DNA damage, ultimately leading to failure in mitosis. Treatment with any of three structurally distinct Chk1/2 inhibitors resulted in increased γH2AX staining in M229R cells over M229P cells, while there was no difference between M229P/R cells under basal conditions (Fig. 4C and D). The increased DNA damage is probably not due to elevated ROS levels, since basal ROS was not elevated in M229R cells (Fig. S3). Overall, these data suggest that Chk1/2 inhibitors selectively induce the accumulation of DNA damage in M229R cells, ultimately leading to a high rate of mitotic failure. This accumulation of DNA damage is likely not a direct effect of Chk1/2 inhibitors, but rather inhibition of Chk1/2 in these cells prevents the repair of DNA damage which is introduced from another source. This, in turn, suggests that the M229R cells may be more prone to DNA damage.

**Figure 4.**
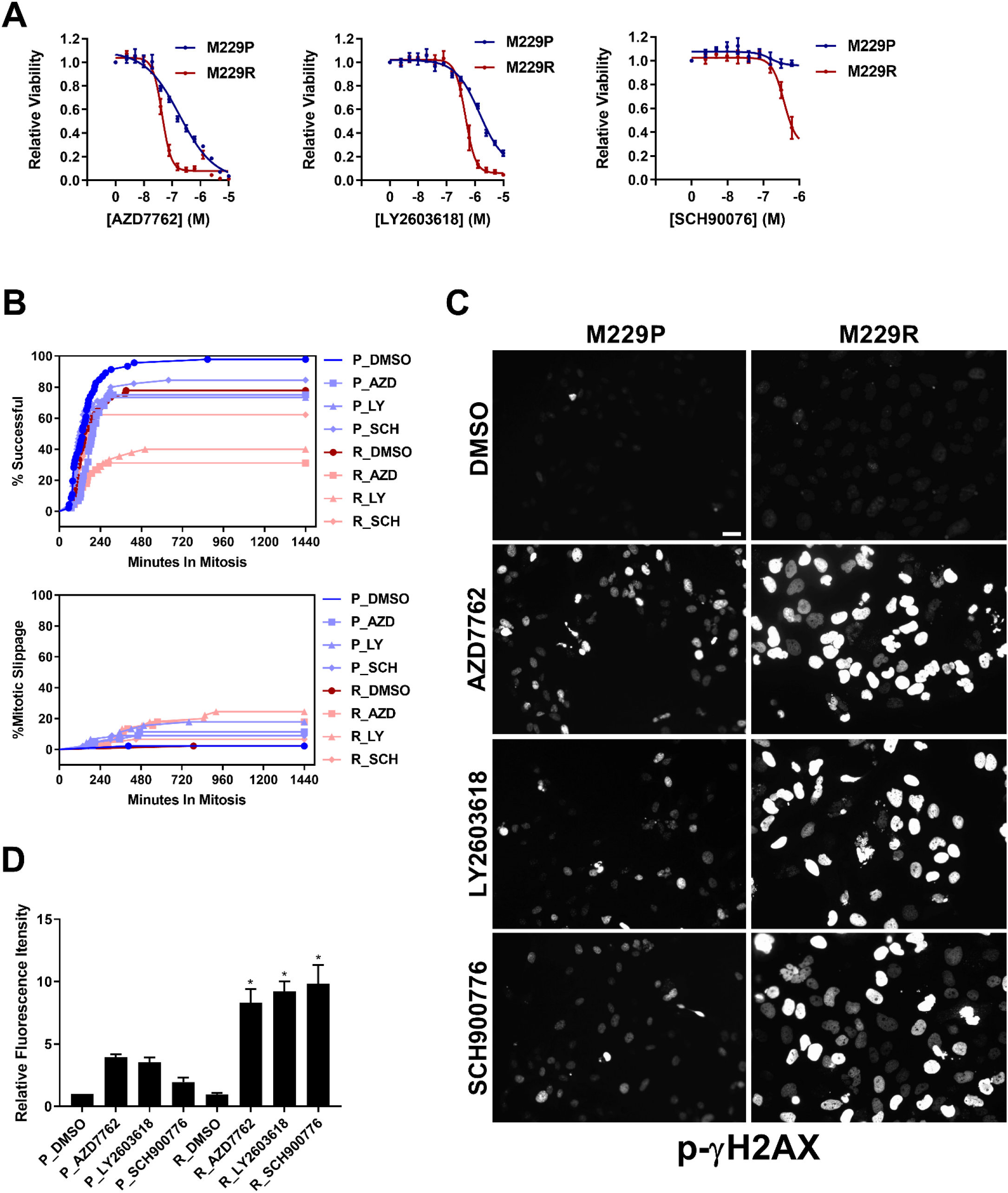
M229R cells are vulnerable to Chk1/2 inhibitors. **A**. M229P/R cells were seeded into 384-well plates and treated with AZD7762, LY2603618, and SCH900776 as indicated. After 72 h, viability was measured as described in Materials and Methods. Data are represented as mean ± SE of the technical replicate averages for each of the biological replicates (n = 3). IC_50_ and E_max_ values are listed in Table S3. **B**. M229P/R cells were engineered to express mScarlet-H2A and EGFP-TUBA1B as described in the Materials and Methods. Cells were seeded into glass-bottom 96-well plates and the next day the cells were treated with 100 nM AZD7762, 1 µM LY2603618, or 1 µM SCH900776. Mitotic rate/outcome was measured on the Cytation 3 microscope setup as described in Materials and Methods. At least 40 cells were analyzed per treatment condition. **C**. M229P/R cells were treated with 100 nM AZD7762, 1 µM LY2603618, or 1 µM SCH900776 for 24 h. The cells were subsequently fixed and stained with an antibody raised against p-γH2AX. Scale bar = 10 µM. **D**. Quantification of γH2AX from the experiment in Figure. 4C was as described in Materials and Methods. Statistical analysis was performed with one-way ANOVA analysis, * indicates p < 0.01 vs the M229R DMSO group. None of the compound-treated M229P groups were statistically significant in comparison to M229P DMSO. Data are represented as mean ± SE for n = 3 biological replicates. IC_50_ and E_max_ values are listed in (Table S3).

## Discussion

In this study through screening of a drug library that BRAFi resistance in examined melanoma cell lines is accompanied by increased sensitivity to either a broad class of mitotic disrupters, including AURK, PLK, tubulin, and kinesin inhibitors or, in the case of one resistant line, to Chk1/2 inhibitors. For the group of BRAFi resistant lines that are rendered sensitive to mitotic inhibitors, our data suggest that the mechanistic basis of this selectivity is an inability of the BRAFi-resistant cells to undergo mitotic slippage. Mitotic slippage is a well characterized resistance mechanism for multiple classes of mitotic inhibitors, including those which disrupt tubulin polymerization/depolymerization (*30-32*). Our data support the idea that the inability of UACC62R cells to exit from mitotic arrest through mitotic slippage upon treatment with the mitotic inhibitor, MLN8237, results from differential Cyclin B1 degradation, since Cyclin B1 degradation is a key initiating event during mitotic slippage. Under physiological conditions, Cyclin B1 is targeted for degradation by the anaphase-promoting complex (APC) during metaphase(*33*). UACC62P/R cells upon treatment with the AURKi MLN8237 appeared to arrest in prophase/prometaphase with condensed chromosomes but no alignment of the chromosomes along the metaphase plate. These data would suggest that the APC is still inactivated in these cells, which should prevent the degradation of Cyclin B1(*34*). It is possible that a low level of APC activation is present in UACC62P, but not UACC62R, cells, which would result in the gradual degradation of Cyclin B1 and eventual mitotic slippage. Another possibility is that the APC may be fully inactivated in both UACC62P and UACC62R cells, but APC-independent Cyclin B1 degradation mechanisms could have higher activity levels UACC62P cells. Further clarification of these mechanisms will be important since they could serve as biomarkers for identifying tumors which are more responsive to disruption of mitosis.

Another BRAFi-resistant cellular model, M229R, acquired increased sensitivity to Chk1/2 inhibitors. While the molecular mechanism governing this selectivity is different from that of UACC62P/R cells, the commonality is that both cellular models are vulnerable to inhibitors which directly or indirectly disrupt mitosis. Chk1/2 inhibitors induced a more severe accumulation of the DNA damage marker γH2AX in M229R cells than in M229P cells. This could indicate that excessive DNA damage is causing the M229R cells to arrest and ultimately die during mitosis. One possible explanation for the differential response to Chk1/2 inhibitors is functional redundancy between Chk1/2 and other DNA repair pathways. For instance, if M229R cells are defective in other DNA repair mechanisms, this would increase their dependence on Chk1/2 for DNA repair, ultimately resulting in an elevated accumulation of DNA damage in Chk1/2i-treated M229R cells. This model would also explain why there is no difference in γH2AX staining in DMSO-treated M229R cells, since in the absence of Chk1/2 inhibitors M229R cells would still retain the ability to perform DNA repair. An analogous model explains why BRCA-mutant tumors have elevated sensitivity to PARP inhibitors (*35*).

In summary, using a drug repurposing screening approach, we observed acquired pharmacological vulnerability to compounds that result in mitotic disruption in three different poorly differentiated BRAFi resistant melanoma cell lines. In contrast, no compound class showed selective toxicity in a cell line known to develop BRAFi resistance through acquisition an NRAS mutation. This observation suggests that melanoma cells and tumors whose resistance is associated with a dedifferentiation phenotype are more vulnerable to compounds which disrupt mitosis. If biomarkers for response to anti-mitotic agents can be established, it may be possible to identify a subset of resistant tumors which are vulnerable to second-line therapy with these classes of approved drugs. While we did not observe synergy between BRAF inhibitors and mitotic inhibitors in BRAFi-resistant cells, the combination of these agents still warrants further investigation. The finding that BRAFi-resistant cells are more sensitive to mitotic inhibitors suggests that tumors may be more sensitive to these agents after they develop resistance to MAPKi therapy. However, another intriguing possibility is that BRAF and mitotic inhibitors could be combined at the onset of treatment to prevent or forestall the development of drug resistance. This is especially true if mechanisms of resistance to BRAF/MEK inhibitors are mutually exclusive to mechanisms of resistance for mitosis inhibitors.

## Acknowledgements

J.C.S is a Damon Runyon Dale F. Frey Scientist supported (in part) by the Damon Runyon Cancer Research Foundation (DFS-24-17). The NCATS MIPE screening library was provided by NCATS.

