## Supplemental Figures for "BRAF inhibitor resistance confers increased sensitivity to mitotic inhibitors"

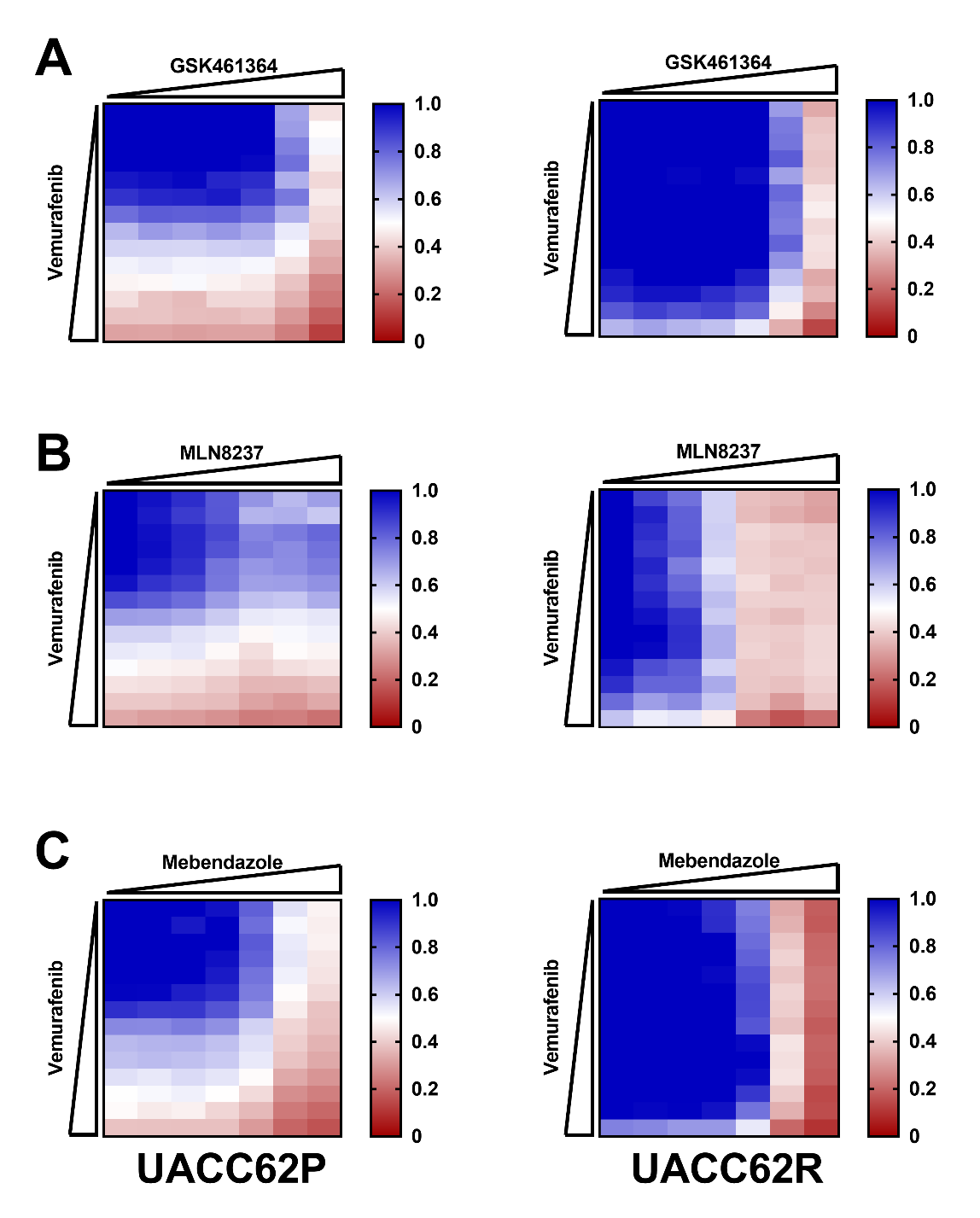

**Figure S1. AURK, PLK, and Tubulin inhibitors do not synergize with vemurafenib in UACC62P/R cells.** UACC62P/R cells were seeded into 384-well plates at a density of 1,000 cells/well. The next day the cells were treated in a concentration response matrix with a top concentration of 10 µM for all compounds and a ½ dilution series. Viability was analyzed as described in the Materials and Methods section. Data is expressed as relative viability wherein a value of 1 (blue) indicates 100% viability and a value of 0 (red) indicates 0% viability. **A.** UACC62P/R cells were treated with a GSK461364 x Vemurafenib concentration response matrix. **B.** UACC62P/R cells were treated with a MLN8237 x Vemurafenib concentration response matrix. **C.** UACC62P/R cells were treated with a Mebendazole x Vemurafenib concentration response matrix. This experiment was repeated with n = 3 biological replicates. The data in these matrices represent the average compound response across all experimental replicates.

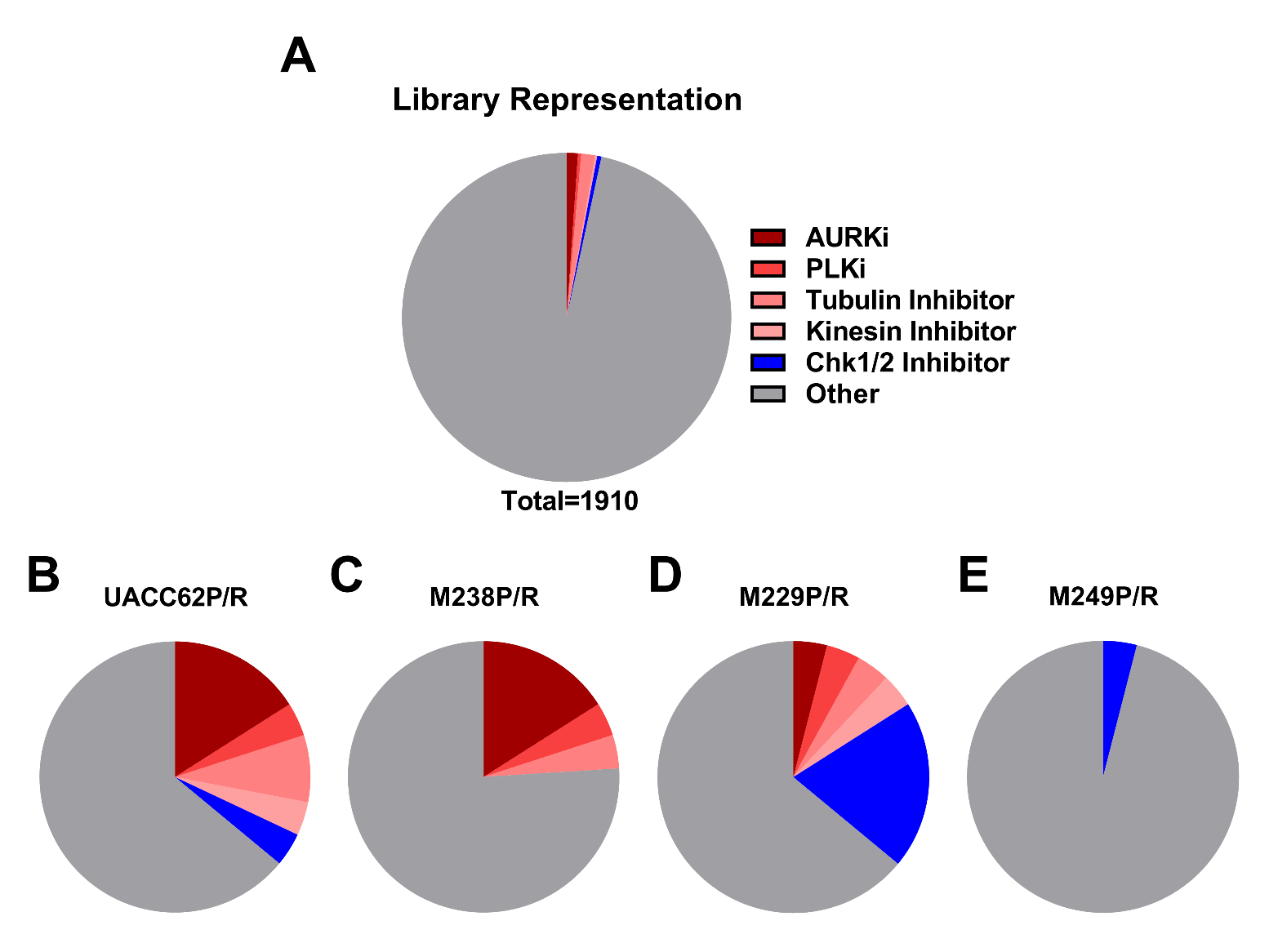
**Figure S2. Identification of compound classes which are selective for BRAFi-resistant cells. A)** Overall compound representation in the MIPE library. AURKi, PLKi, Tubulin inhibitors, Kinesin inhibitors, and Chk1/2 inhibitors are highlighted. Compound class enrichment for the top 25 most selective compounds in **B)** UACC62P/R cells, **C)** M238P/R cells, **D)** M229P/R cells, and **E)** M249P/R cells.

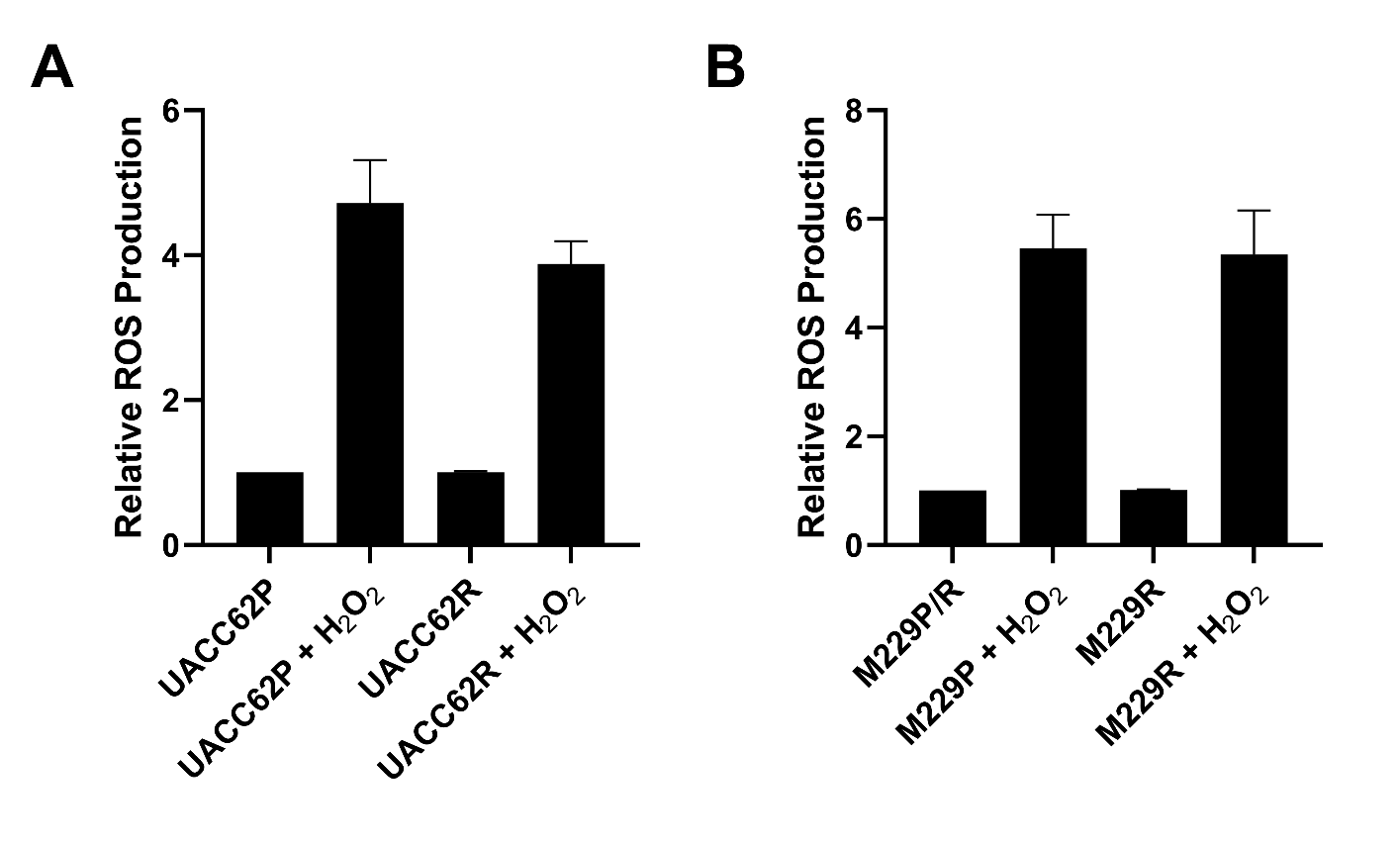

**Figure S3. ROS production is not altered in BRAFi-resistant cells.** **A)** UACC62P/R and **B)** M229P/R cells were seeded into 96-well plates. The next day the cells were treated with H_2_O_2_ and the ROS assay was performed as described in the Materials and Methods with cells treated with H_2_O_2_ serving as a positive control. Relative ROS production is normalized to ROS levels in UACC62P (panel A) or M229P (panel B). This experiment was repeated with n = 3 technical replicates and n = 3 biological replicates.

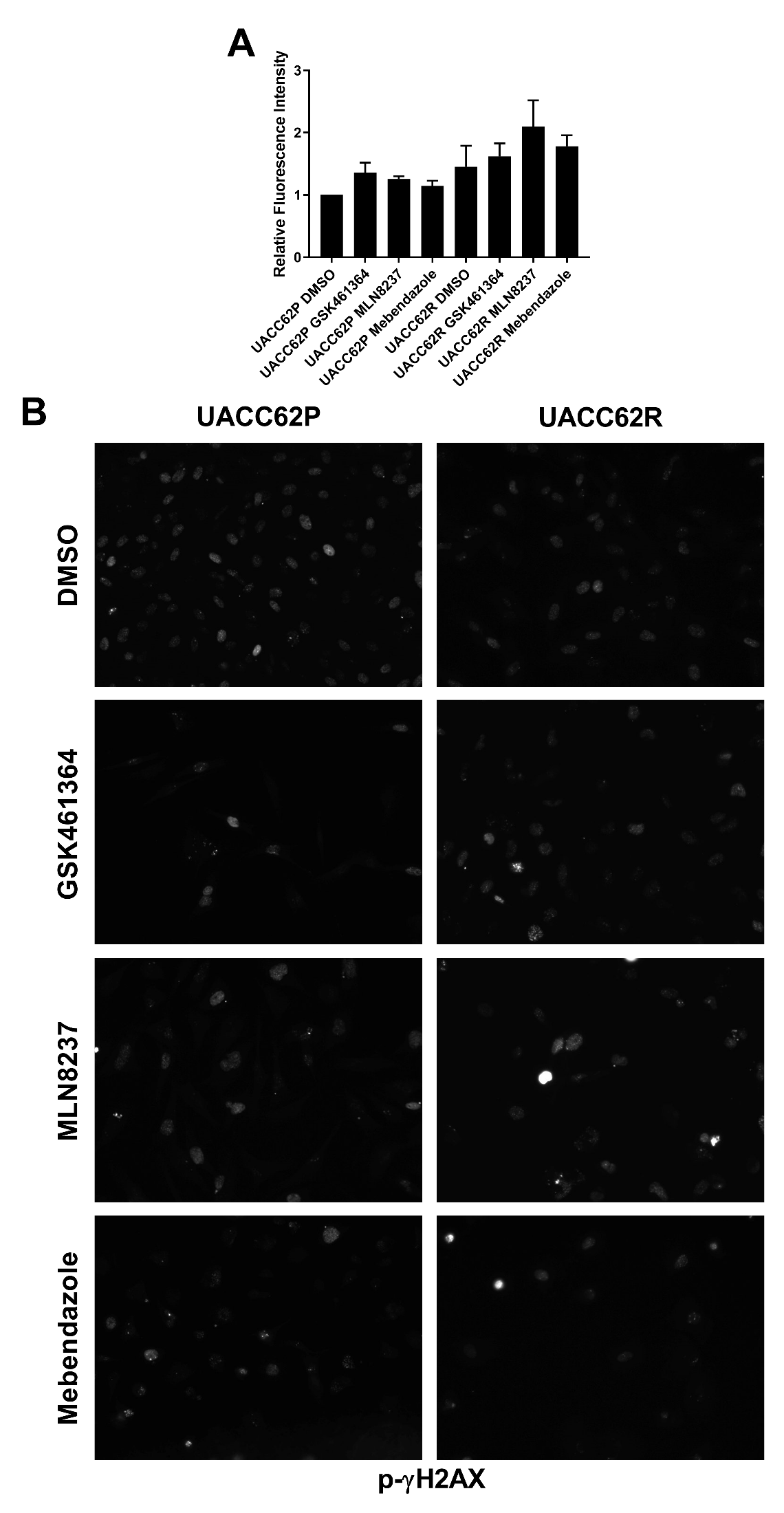

**Figure S4. p-γH2AX staining is not altered in compound-treated UACC62P/R cells. A**) UACC62P/R cells were treated with 1 µM GSK461364, MLN8237, or Mebendazole for 24 h. The cells were fixed and stained with a p-γH2AX antibody and quantified as described *Materials and methods*. **B)** Representative immunofluorescence images. This experiment was repeated with n = 3 biological replicates.

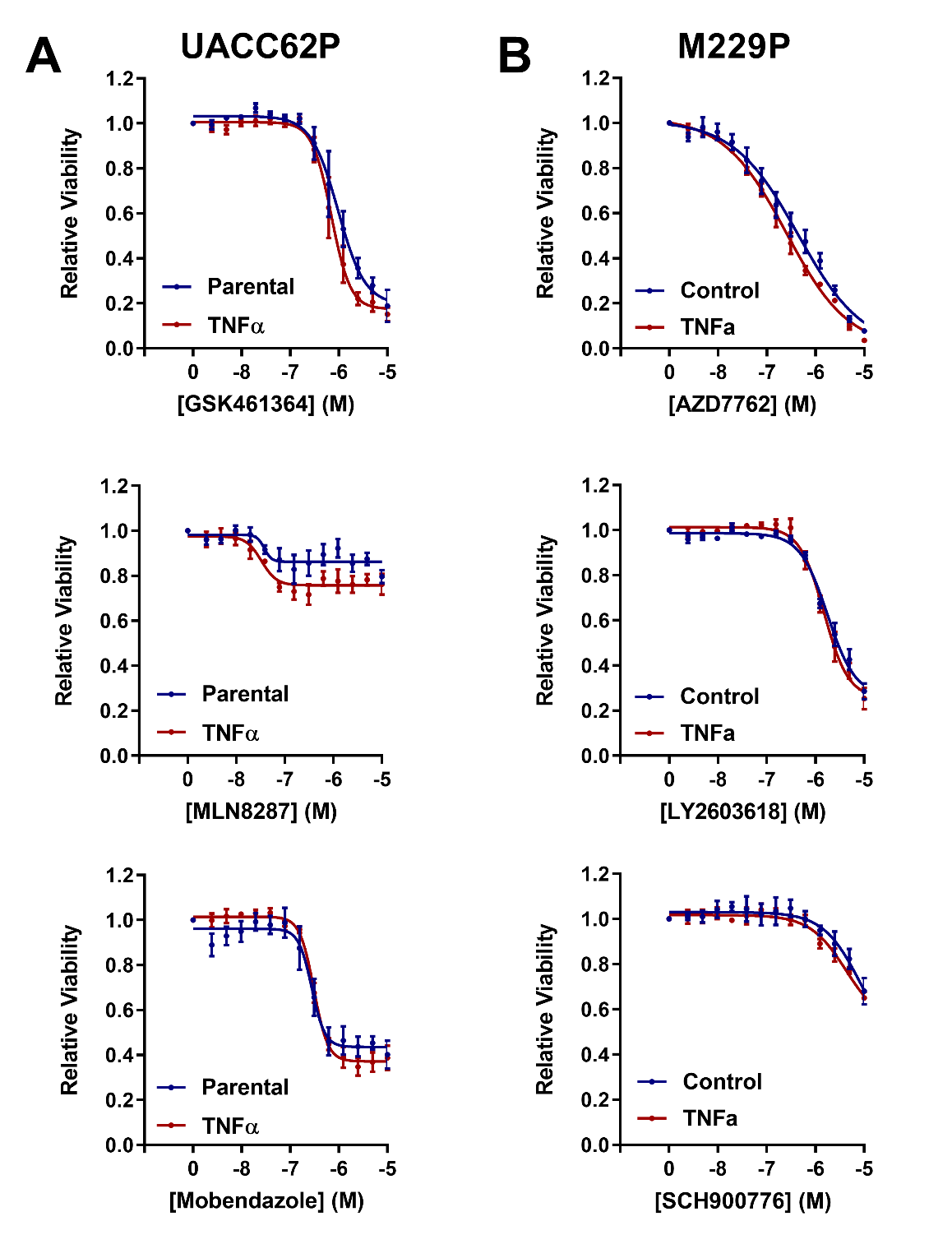

**Figure S5. TNFα does not alter AURK, PLK, Tubulin, or Chk1/2 inhibitor sensitivity. A)** UACC62P or **B)** M229P cells were seeded into 384-well plates at a density of 1,000 cells/well. The next day the cells were treated -/+ 10 ng/mL TNFα and a concentration gradient of the indicated compound. Viability was measured and quantified as described in the Materials and Methods section. This experiment was repeated with n = 3 technical replicates and n = 3 biological replicates.

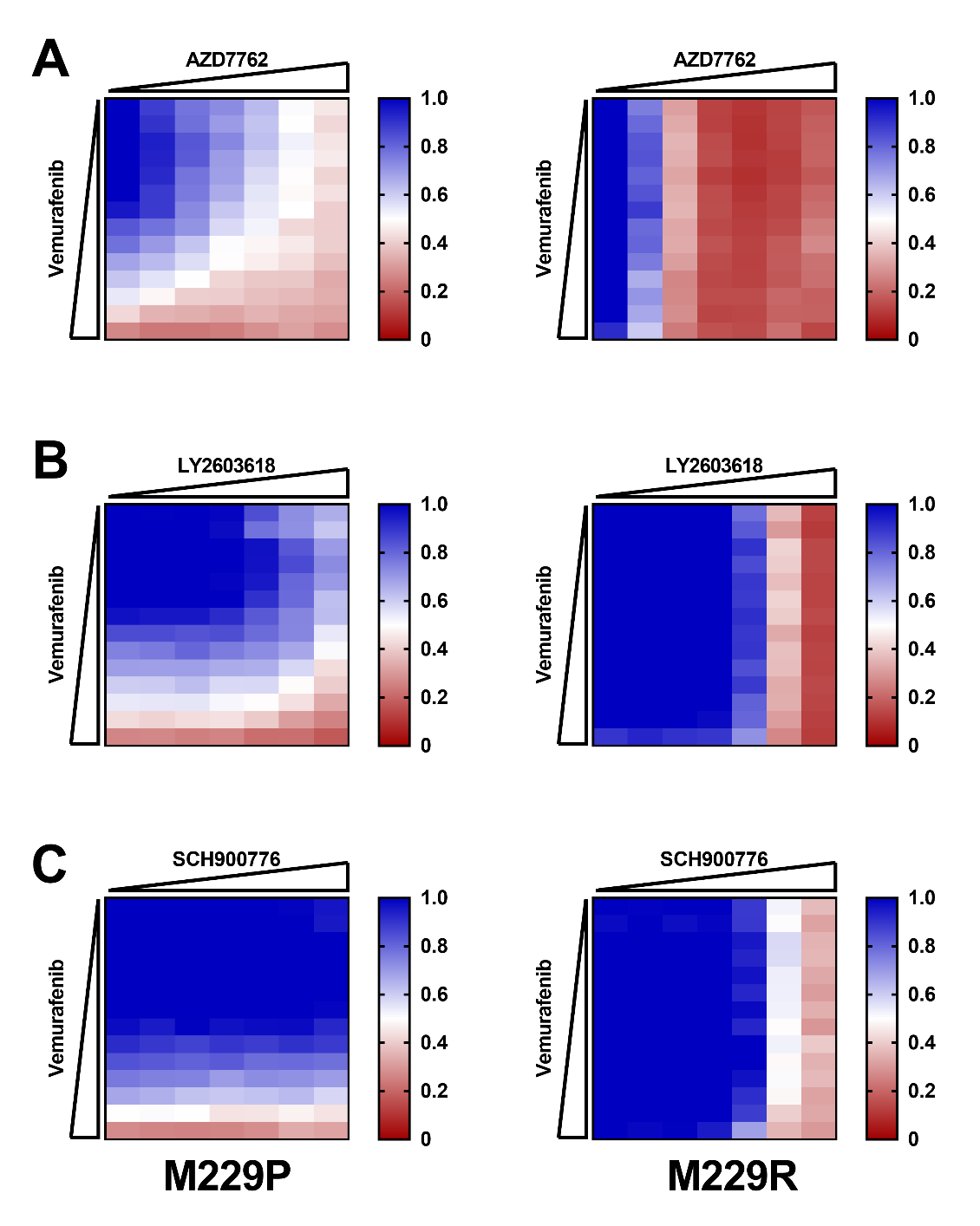

**Figure S6. Chk1/2 inhibitors do not synergize with vemurafenib in M229P/R cells.** M229P/R cells were seeded into 384-well plates at a density of 1,000 cells/well. The next day the cells were treated in a concentration response matrix with a top concentration of 10 µM for all compounds and a ½ dilution series. Viability was analyzed as described in the Materials and Methods section. Data is expressed as relative viability wherein a value of 1 (blue) indicates 100% viability and a value of 0 (red) indicates 0% viability. **A.** M229P/R cells were treated with a AZD7762 x Vemurafenib concentration response matrix. **B.** M229P/R cells were treated with a LY2603618 x Vemurafenib concentration response matrix. **C.** M229P/R cells were treated with a SCH900776 x Vemurafenib concentration response matrix. This experiment was repeated with n = 3 biological replicates.
